# Impact of environmental factors on calling behavior in midshipman fish across ocean basins

**DOI:** 10.1101/2023.07.11.547287

**Authors:** Annebelle C.M. Kok, Ella B. Kim, Timothy J. Rowell, Tetyana Margolina, John E. Joseph, Lindsey E. Peavey Reeves, Leila T. Hatch, Simone Baumann-Pickering

**Affiliations:** Acoustic Ecology Laboratory, Scripps Institution of Oceanography, University of California, San Diego, La Jolla, CA 92037, USA; Northeast Fisheries Science Center, National Oceanic and Atmospheric Administration, Woods Hole, MA 02543, USA; Oceanography Department, Naval Postgraduate School, Monterey, CA 93943, USA; National Marine Sanctuary Foundation, Silver Spring, MD 20910, USA; Office of National Marine Sanctuaries, National Oceanic and Atmospheric Administration, Silver Spring, MD 20910 USA

## Abstract

Chorusing is widespread across the animal kingdom. Animal calling behavior is often driven by phenological and environmental factors such as seasonality, lunar period, and temperature. Now, in the Anthropocene, factors such as increased anthropogenic noise levels are also affecting calling behavior. Many fish call in choruses to attract mates, but the dynamics that drive fish calling behavior have rarely been studied in the field. We investigated how seasonality, lunar period, ambient noise, and temperature influenced the calling behavior of two species of toadfish, the plainfin midshipman (*Porichthys notatus*) and putatively, the Atlantic midshipman (*Porichthys plectrodon*). Acoustic recordings from a two-year period in twelve different locations, spanning two ocean basins showed that midshipman chorus presence was driven by seasonality and lunar period. Furthermore, chorus frequency increased with increasing temperature. Chorus levels were strongly influenced by seasonality and increased somewhat with increasing noise levels. Taken together, these results indicate that midshipman calling behavior was strongly influenced by interacting environmental conditions. Understanding the various impacts of each driver will facilitate predictions of changes in midshipman calling due to future changes in environmental conditions.

## INTRODUCTION

Many animals time their signaling activity at a specific period of the day. Songbirds call predominantly in the morning, during the dawn chorus, with a species-specific peak chorusing time (Thomas et al. 2002; Bruni et al. 2014). Frogs often chorus in the evening and form densely aggregated chorusing groups to attract mates (Stratman et al. 2021). In fish, chorusing activity can be so intense that it dominates the marine soundscape (McKenna et al. 2021). The presence of conspecific calls can increase calling in individuals (Bee and Perrill 1996; Capshaw et al. 2020), and environmental factors often drive the temporal patterns in calling activity (Bruni et al. 2014; Ladich 2019).

In fish, birds, and frogs, one function of calling is to attract mates and repel same-sex rivals (Lobel 1992; Gerhardt 1994; Catchpole and Slater 2003). For many bird and frog species, this link between calling activity and the mating season is well-established (Dawson et al. 2001). For fish, less information is available but to date, evidence points toward a similar synchrony between calling activity and spawning. For example, goliath groupers (*Epinephelus itajara*) produced ‘boom’ calls on nights when spawning was confirmed by egg collection, but did not call on nights without egg deposits (Koenig et al. 2017). Similarly, plainfin midshipman (*Porichthys notatus*) males attracted females to their nests with humming sounds, where the females spawned (Brantley and Bass 1994).

Another factor that influences calling activity in both terrestrial and marine animals is the lunar cycle (Staaterman et al. 2014). Some fish species call more during new moon nights, and cease activity during full moon nights (Parsons et al. 2016; Kaplan et al. 2018), while others increase acoustic activity during full moon nights (Stanley et al. 2020; Borie-Mojica et al. 2022) or other specific lunar periods, which can be site or species specific (Mann et al. 2010; Rowell et al. 2015). In birds, the lunar cycle affects chorus onset time: birds called earlier on mornings when the moon had been in the third quarter or full and was still present above the horizon at dawn (Bruni et al. 2014). In both fish and birds, lunar illumination is the driving factor. Light presence increases the possibility to detect prey or to be detected by predators, which both come as a trade-off to calling for mates (Thomas et al. 2002; Maiditsch and Ladich 2022).

An environmental factor that drives calling activity especially in ectotherms is temperature. In fish, increased temperature often leads to increased calling activity or an increase in the peak frequency of the call (Ladich 2018). However, these patterns have only conclusively been found in the lab. In the field, studies found contradicting results, which is possibly related to other environmental factors interacting with the effect of temperature, such as seasonality (Ladich 2018). Unraveling the true effect of temperature on fish calling in the field will therefore require sampling across one or multiple seasons at multiple locations to correct for these interacting effects.

Finally, ambient noise levels can also influence calling behavior in terrestrial and marine animals. When ambient noise levels increase, animals adapt their calling behavior in a variety of ways: they may increase their call amplitude or repetition rate to compensate for the masking effect of the noise, known as the Lombard effect (Brumm and Zollinger 2011); extend call duration (Bermúdez-Cuamatzin et al. 2011); shift the frequency of their calls to be outside of the frequency range of the noise (Roca et al. 2016); or cease calling when noise levels are too high (Tsujii et al. 2018). However, the compensation is usually not equal to the increase in noise level, leading to a lower signal-to-noise ratio (Guazzo et al. 2020). Although not all animals exhibit the Lombard effect (e.g. Cope’s grey treefrog, *Hyla chrysoscelis* (Love and Bee 2010)), the occurrence of the Lombard effect has been shown across a wide range of taxa. In fish, the Lombard effect was described for multiple species, including the plainfin midshipman (Holt and Johnston 2015; Luczkovich et al. 2016; Brown et al. 2021). In an experimental setup, nest-guarding midshipman fish were exposed to artificially increased noise levels. Although the overall trend was an increase in call levels, the variation in call levels also increased significantly, indicating that not all individuals followed the same strategy.

To understand how these environmental drivers operate on the calling behavior of both marine and terrestrial animals, it is important to look for patterns that transcend a single location and a single species. Therefore, we studied environmental drivers on the chorusing behavior of two marine fishes that produce similar calls: the chorus from plainfin midshipman and the chorus that is presumed to be from Atlantic midshipman (*Porichthys plectrodon*). The species both occur in shallow waters along the coast of North America: plainfin midshipman in the Pacific ocean, Atlantic midshipman in the Atlantic ocean. Both choruses consist of harmonically structured humming sounds (McIver et al., 2014, Fig. 1). The choruses can last for several hours, without obvious breaks and can dominate discrete frequencies of the soundscape (McKenna et al. 2021). The fundamental frequency of presumed Atlantic midshipman hums is typically higher than that of the plainfin midshipman, 180 Hz vs 100 Hz, respectively. We investigated 1) the temporal patterns of midshipman chorusing, and 2) the influence of environmental drivers – lunar cycle, temperature and ambient noise – on chorus presence, chorus level and peak frequency of the chorus. By comparing ambient conditions across sites, we hope to elucidate potential changes in the calling behavior of fish in a variable marine environment.

**Figure 1:**
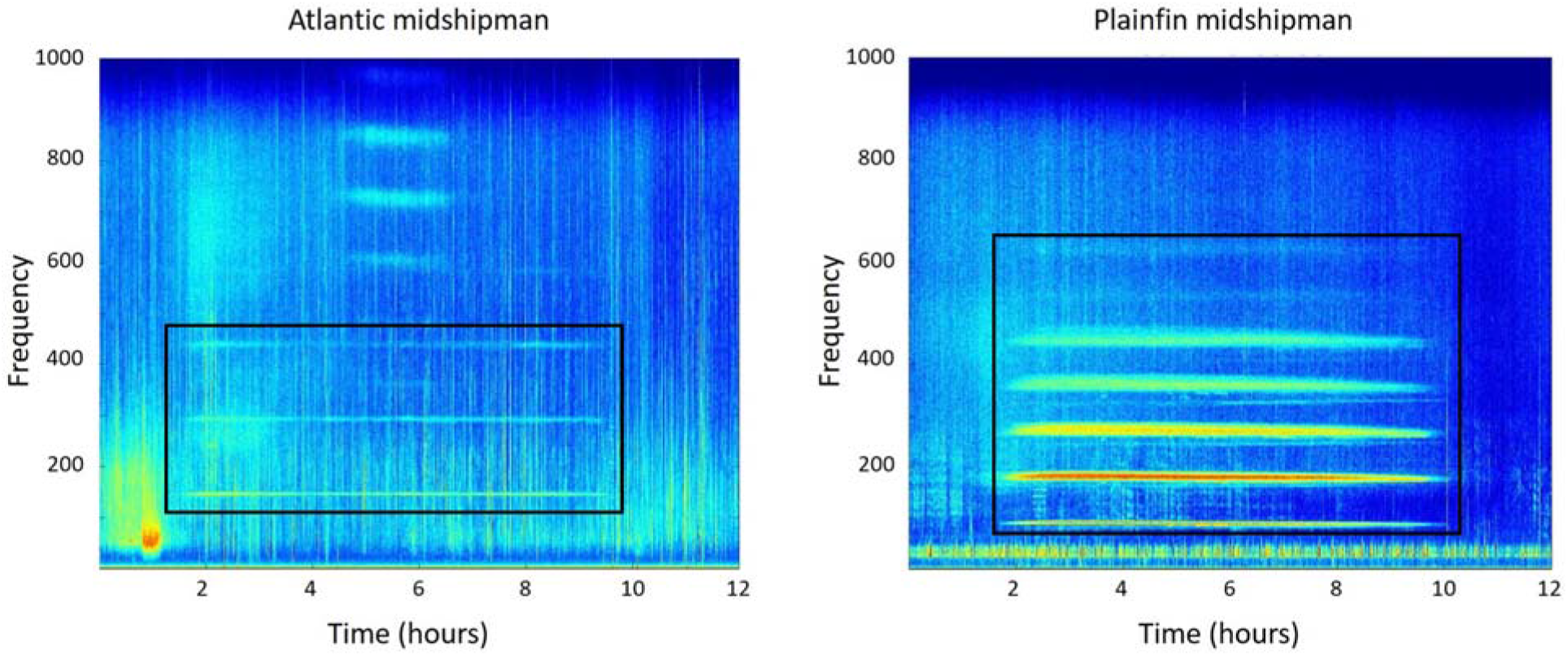
Long-term spectral average of a putative Atlantic midshipman hum (left) and plainfin midshipman hum (right). Choruses are marked by black boxes. Long-term spectral averages were calculated in 5 s, 1 Hz bins. Note the presence of other fish choruses on the panel with the Atlantic midshipman hum, that are obscuring the left part of the boxed area and are also visible outside the boxed area.

## METHODS

### Study sites

This study was part of a larger collaboration, called SanctSound. The U.S. National Oceanic and Atmospheric Administration (NOAA) and the U.S. Navy engaged in a multi-year effort (2018-2022) to monitor underwater sound within the U.S. National Marine Sanctuary System. The study included recording stations in sanctuaries off the east coast of the U.S. (Stellwagen Bank, Gray’s Reef, and Florida Keys National Marine Sanctuaries), the west coast of the U.S. (Olympic Coast, Monterey Bay, and Channel Islands National Marine Sanctuaries), and the Pacific region (Hawaiian Islands Humpback Whale National Marine Sanctuary and Papahânaumokuâkea Marine National Monument).

SanctSound was designed to provide standardized acoustic data collection and analysis to document how much specific sources contributed to the soundscape present within these protected areas as well as potential impacts of noise to the areas’ marine taxa and habitats. To understand the features of a given listening station that influence measured sound levels, the SanctSound effort encompassed results from multiple acoustic detection algorithms (biological and anthropogenic), data from a wide range of non-acoustic variables (e.g. gliders, vessel traffic data, weather stations), and sound propagation models to quantify variation in specific sound source detection ranges.

This analysis assessed midshipman chorusing at 12 SanctSound recording sites within four National Marine Sanctuaries: Gray’s Reef NMS and Florida Keys NMS in the western Atlantic, and Channel Islands and Monterey Bay NMSs in the eastern Pacific (Fig. 2; Table 1). No midshipman chorusing was found at the other National Marine Sanctuaries (Olympic Coast NMS, Stellwagen Bank NMS, Hawaiian Islands Humpback Whale NMS and Papahânaumokuâkea Marine National Monument).

**Table 1:** Site information of recording locations in the Atlantic (N=4) and Pacific (N=8).

| Region | Sanctuary | Site ID | Location | Water depth (m) |
| --- | --- | --- | --- | --- |
| Atlantic | Florida Keys | FK01 | 24.433130 N 81.930680 W | 15 |
|  | Florida Keys | FK02 | 24.488800 N 81.666316 W | 13 |
|  | Florida Keys | FK03 | 24.458950 N 81.774133 W | 25 |
|  | Gray's Reef | GR01 | 31.396417 N 80.8807 W | 17 |
| Pacific | Channel Islands | CI01 | 34.0438 N 120.0801 W | 21 |
|  | Channel Islands | CI02 | 34.0856 N 120.5232 W | 77 |
|  | Channel Islands | CI03 | 33.48687 N 119.0151 W | 29.5 |
|  | Channel Islands | CI04 | 33.849 N 120.118 W | 156 |
|  | Channel Islands | CI05 | 34.0178 N 119.3172 W | 139 |
|  | Monterey Bay | MB01 | 36.798 N 121.976 W | 120 |
|  | Monterey Bay | MB02 | 36.6495 N 121.9084 W | 68 |
|  | Monterey Bay | MB03 | 36.37021 N 122.314903 W | 858 |

**Figure 2:**
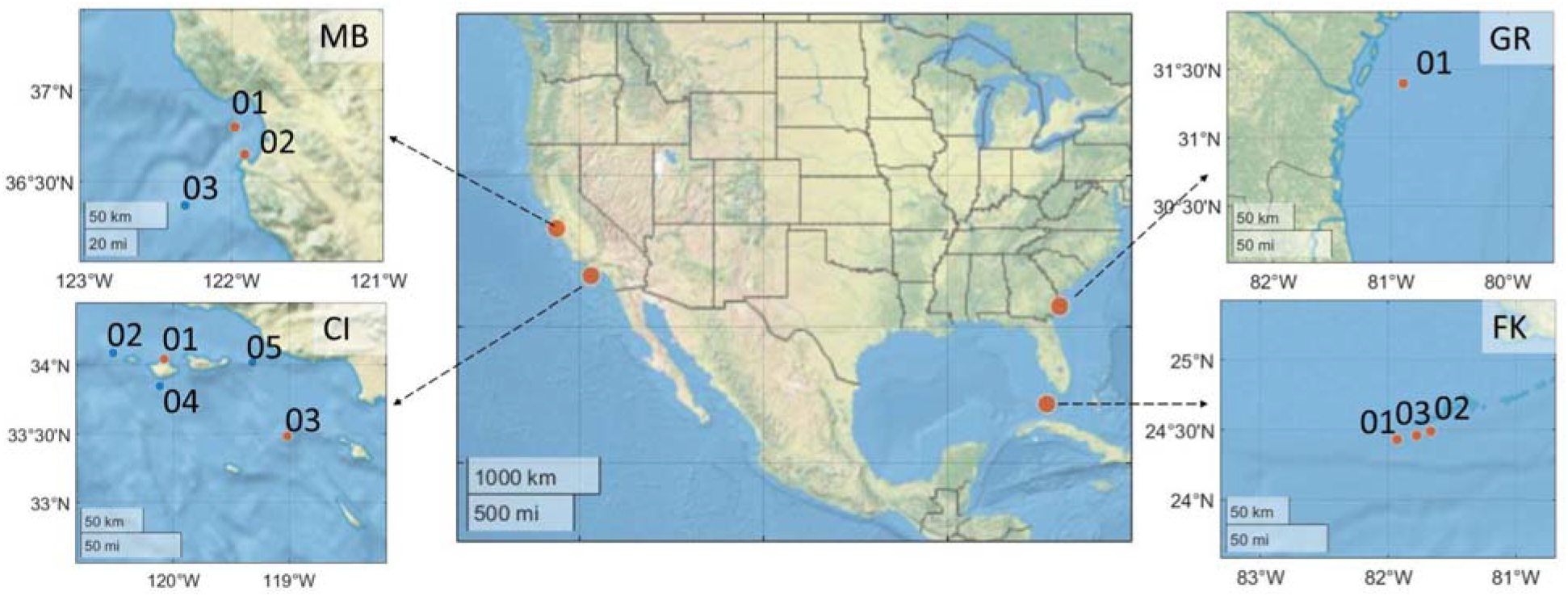
Midshipman chorusing was recorded at most (orange) but not all (blue) recording locations. Locations were numbered within sanctuaries: MB = Monterey Bay; CI = Channel Islands; GR = Gray’s Reef; FK = Florida Keys.

Five SoundTrap (Ocean Instruments, NZ) recorders (version 500STD) were placed at Channel Island National Marine Sanctuary (CI01-05), three at Monterey Bay National Marine Sanctuary (MB01-02), one recorder at Gray’s Reef National Marine Sanctuary (GR01) and three recorders at Florida Keys National Marine Sanctuary (FK01-03). All recorders except one at Monterey Bay were SoundTrap (Ocean Instruments, NZ) recorders (version 500STD). The third recorder at Monterey Bay was a high-frequency acoustic recording package (Wiggins and Hildebrand 2007). All recorders in the Pacific were moored 1-3.5 m from the ocean floor with a subsurface float and an acoustic release, while the recorders in the Atlantic were attached to fixed platforms at a depth of 1 m above the seafloor and were deployed by divers. The Atlantic sites did not have acoustic releases nor subsurface floats. All but one recorder (MB03 at 858 m) were placed at water depths of 13-156 m.

### Acoustic recordings

Sound was recorded between November 2018 and July 2020, with gaps due to servicing of recorders and some equipment malfunction (Fig. S1). Recording sampling rate was 48 kHz (96 kHz for the first deployment of CI01, CI02, CI04, CI05, MB01, and MB02, 200 kHz for MB03). During analysis, all recordings were decimated to 2 kHz to improve processing speed. The recordings were visualized in long-term spectral averages (LTSAs), with 5-sec and 1-Hz resolution over 12 h scanning windows to capture a full chorusing event per day (Fig 1). Recording data used in this analysis are available via the project data portal (https://data.noaa.gov/waf/NOAA/NESDIS/NGDC/MGG/passive_acoustic//iso/).

### Ancillary data collection

Temperature measurements were taken at 5 s intervals from a temperature sensor on the acoustic recorders for all sites, which were offset from true water temperature by 0.5-1 °C (determined by comparing measurements from the temperature sensors to nearby telemetry receivers at Florida Keys NMS). For Florida Keys NMS (FK), there were gaps in the temperature data which were filled with temperature measurements from a nearby telemetry receiver (∼300 m distance, 24.43070 N, 81.93180 W). For Channel Islands NMS 1 (CI01), temperature data was missing between 09-04-2020 and 02-06-2020, for which there was no nearby temperature measuring station. Erroneous measurements were removed from the data, and data were subsampled to 1 min intervals to match the frequency measurements of the chorus. Lunar illumination data were downloaded from the Horizon server (https://ssd.jpl.nasa.gov/horizons/app.html#/) at a resolution of 30 minute increments. Illumination was recorded in percentages, with 100% representing the full moon and 0% representing the new moon, or no moon present in the sky. For the Channel Islands NMS recordings, wind data was requested from NOAA NDBC Station 46053 (34.241 N, 119.839 W). Wind was reported as average wind speed in m/s every 10 minutes, and was averaged over hourly bins for this study.

### Analysis

The LTSAs were manually scanned for encounters of fish chorusing. In some cases, the exact start or end time could not be determined because it was obscured by noise from other sources. In that case, the first/last visible instance of the chorus was logged as the start/end time. Next, we used a custom-written script (MATLAB 2016b, Mathworks, Natick, MA) to automate measurements of the chorus peak frequency for each 5-sec bin. To get an accurate automated measurement of the peak frequency, we restricted the search for a maximum received level to a 100 Hz bandwidth around the suspected peak frequency of the chorus. For the plainfin midshipman, this band was in the 2nd harmonic of the chorus at 150-230 Hz. For putative Atlantic midshipman, the strongest harmonic was the fundamental frequency. This harmonic varied throughout the season due to the change in peak frequency of the chorus over time. Therefore, we chose a band of 50-150 Hz at the start of the season and 80-180 Hz from the middle to end of the season when chorus peak frequency was highest. The 5-sec measurements were summarized to 1-min median values.

#### Accounting for background noise levels

When measuring the level of a sound, one measures both the pressure of the sound itself as well as the underlying noise at the same frequency and time. Because of this, sounds with low received levels can be artificially elevated if they are measured in high noise conditions, and hence are unreliable measurements. Removing these measurements from the dataset is typically done by investigating the signal-to-noise ratio. This is calculated by subtracting a measure of the noise level close in time and at the same frequency of the sound from a measure of the signal level. Measurements with a signal-to-noise ratio of 10 or more get >=90% of their pressure amplitude from the signal. Therefore, they are considered reliable measurements.

Because the midshipman chorus lasted several hours at a time, it was not possible to measure a noise level close in time with the same frequency. As an alternative, we measured the noise level at a different frequency, at the same time of the measurement of the chorus. To investigate the validity of our method, we investigated how increased noise levels would affect chorus level measurements by doing a sensitivity analysis of simulated increased noise levels on choruses in low noise conditions.

For the simulation data set, we selected chorus hours with a median noise level of 70 dB or lower and measured the peak frequency of the chorus every minute. Next, we simulated noisy conditions by artificially adding noise from another part of the data to the recordings. We then remeasured the chorus levels and calculated the difference in chorus level between the two measurements. Any chorus level measurements in the simulation data set that increased by 2 dB or more in the noisy conditions were considered to be contaminated (Fig S2). The noise simulations were used to indicate how reliable the chorus level measurements would be, as well as to determine the best approach to remove possible erroneous data points (Fig S3).

Based on the simulation data set, we decided to calculate an hourly running average of the chorus peak frequency and removed all values that were outside 1 standard deviation (SD) of the hourly mean. Of the remaining measurements, we only included chorus levels of 70 dB or higher, with a signal-to-noise ratio (SNR) of 10 or higher. Finally, we calculated hourly median values of the chorus levels and chorus peak frequency. This selection method reduced the available range of noise levels from 109 to 91 dB, but did not reduce the median and 3rd quartile of the distribution (63-66 vs 65-69 median-3rd quartile, original vs final data; Fig. S4).

The chorus level for each 1-min bin was measured as the median power spectral density (PSD) at the chorus peak frequency for that bin. Code for calculating power spectral densities from audio recordings are available on GitHub (Github.com/MarineBioAcousticsRC/Triton, Soundscape Metrics Remora). To investigate ambient noise as a driver of chorus levels we measured ambient noise in 1-min bins. We measured ambient noise levels as the average PSD (averaged in dB) over 150-155 Hz. This band was outside of the fish chorus frequencies (155-230 Hz for Pacific, 110-148 Hz for Atlantic) and was a center frequency band of vessel noise. SNR was calculated by subtracting noise levels from the chorus level. We manually excluded periods in which the chorusing was obscured by other sounds and only included time bins with a SNR of 10 or higher to minimize the influence of overlapping sounds on the chorus level measurements.

### Statistical analysis

We first investigated the relationship between lunar illumination and hourly chorus presence per site, where presence was scored as 0 (not present) or 1 (present). Only months with chorus presence were included in this analysis. As seasonality was not included in this model, we did not have to account for temporal autocorrelation and could fit a generalized linear model with binomial distribution. Second, we investigated the influence of seasonality, temperature, lunar illumination and natural ambient noise levels on the peak frequency and level of midshipman choruses. For this analysis, we took median hourly values of chorus peak frequency, chorus level, and ambient noise. Because of the inclusion of seasonality, sample sizes for MB01, FK01, FK02 and FK03 were too low to fit statistical models. Therefore, these sites were excluded from this analysis.

To correct for temporal autocorrelation, we fit the models using Generalized Estimating Equations (GEEs). GEEs correct for temporal autocorrelation by means of a variance-covariance matrix, which specifies the co-variability in the data that is not explained by the model parameters. We ran generalized linear GEEs from the package *geepack* (Højsgaard et al., 2005) with a Gaussian distribution, a first-order autocorrelation with day as a blocking unit and robust standard error for all environmental models. All statistical models were run in RStudio (version 1.2.5003, RStudio Inc.).

## RESULTS

### Spatial and temporal patterns

Midshipman chorusing was recorded at some recording sites in both the Atlantic and the Pacific (Fig. 2). In the Pacific, midshipman chorusing was recorded at 4 out of 8 sites. The sites without midshipman chorusing were deeper on average (77, 139, 156, and 858 m) than sites at which midshipman chorusing was detected (21, 29.5, 68, and 120 m), suggesting that water depth was related to chorus presence. In the Atlantic, midshipman chorusing was detected at all sites in some amount, but most midshipman chorusing was detected in Gray’s Reef NMS (GR01, Fig. 3).

**Figure 3:**
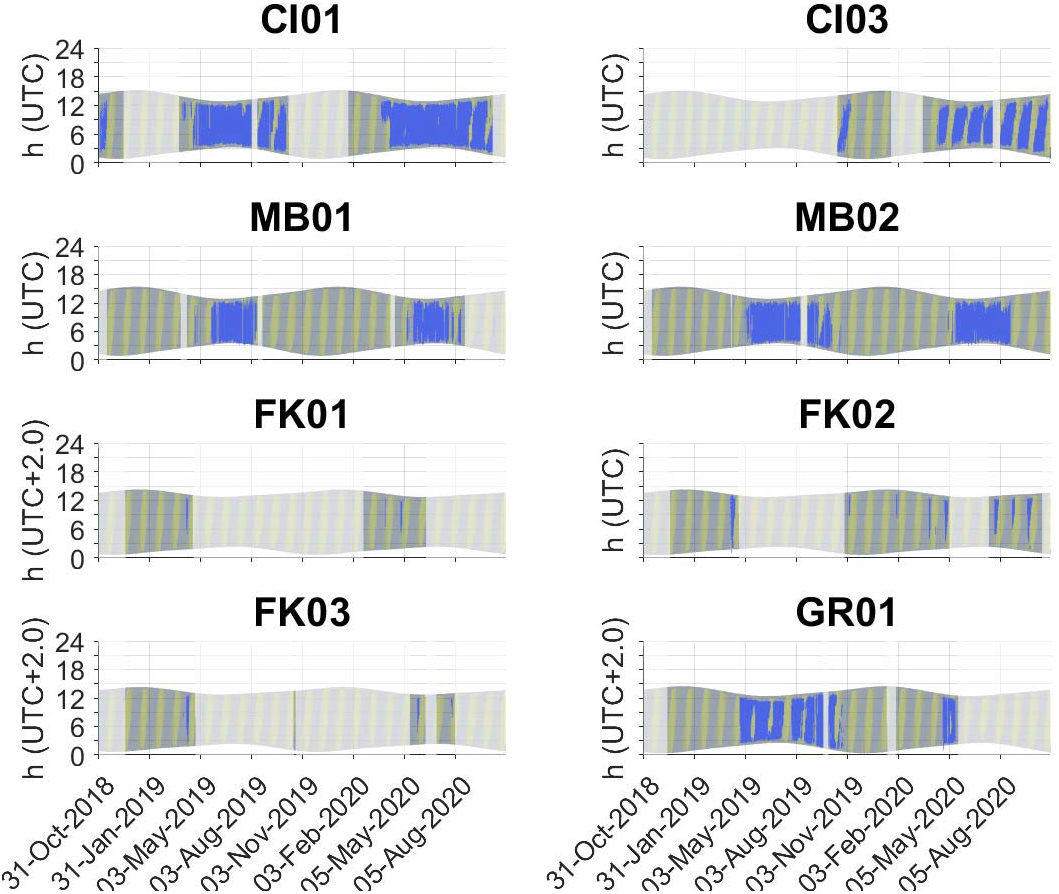
Top panels: temporal patterns of plainfin midshipman chorusing at two sites in Channel Islands National Marine Sanctuary (CI01, CI03) and two sites in Monterey Bay National Marine Sanctuary (MB01, MB02). Bottom panels: temporal patterns of putative Atlantic midshipman chorusing at three sites in Florida Keys National Marine Sanctuary (FK01, FK02, FK03) and one site at Gray’s Reef National Marine Sanctuary (GR01). For both species, chorusing (blue lines) occurred at night (gray shaded areas) and tended to avoid periods with high lunar illumination (yellow shading). The main chorusing period was in late spring and summer. Periods without effort are shaded in white.

In both the Pacific and the Atlantic, midshipman calling commenced in spring, peaked in early summer and gradually declined in late summer or early fall (Fig. 3). The duration of the season varied with latitude: the two more northerly situated sites in Monterey Bay NMS showed a later start date and an earlier end date than around the Channel Islands and Gray’s Reef NMSs, congruent with the shorter summers at higher latitudes. Chorusing was less prevalent in Florida Keys NMS, but sampling at these sites experienced large data gaps during times when chorusing occurred in other regions, so seasonality remains uncertain for these sites. The number of hours with chorus presence was highest in Channel Islands NMS. At all sites except the two sites with chorus presence in Monterey Bay NMS (MB01 and MB02), hourly chorus presence was related to lunar illumination, with a higher likelihood of chorus presence at lower lunar illumination (Fig. 4, Table S1).

**Figure 4:**
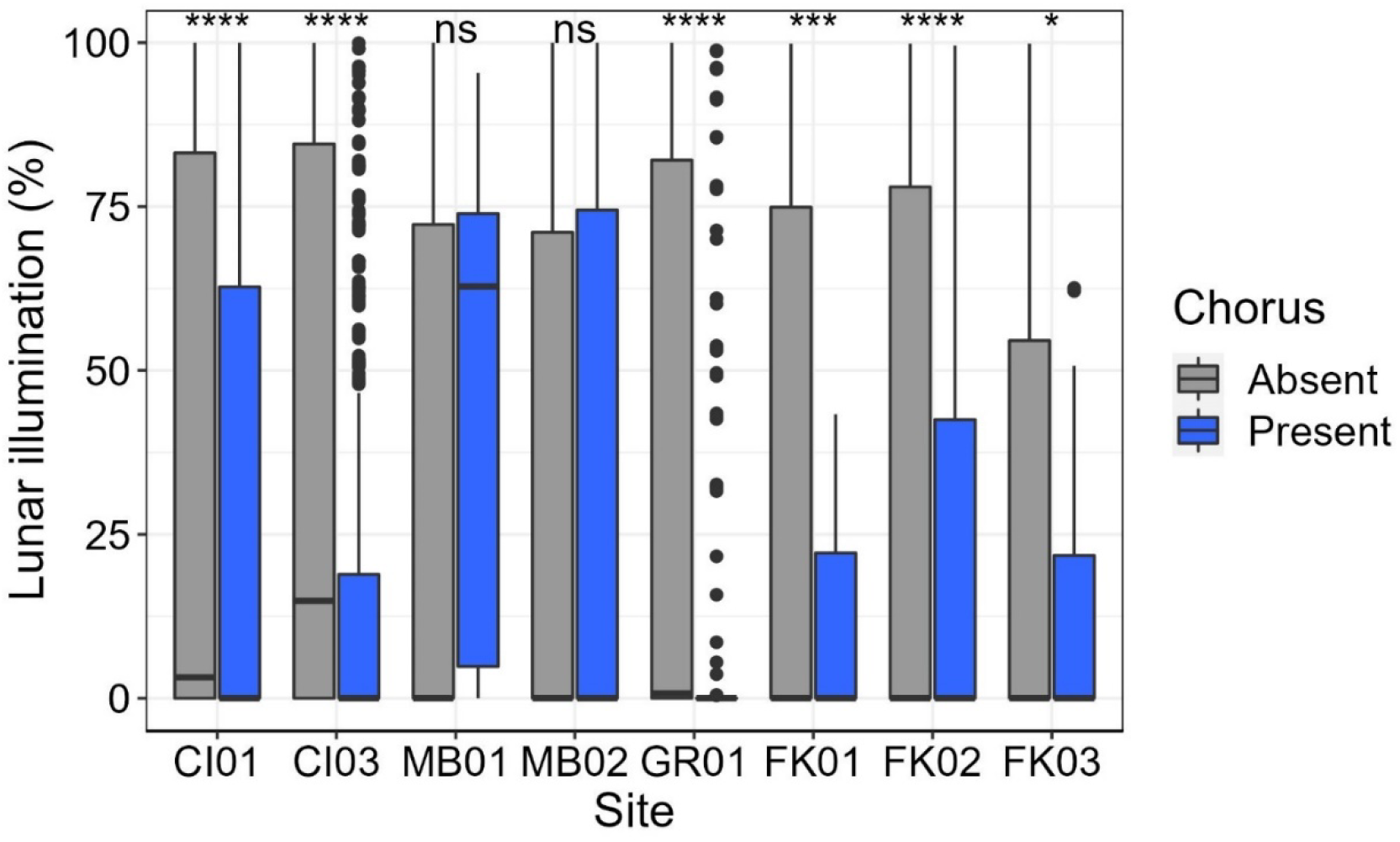
Chorus presence was associated with lower lunar illumination at all sites except MB01 and MB02. Boxes represent 25-75% quantiles, black center lines represent median values per site, whiskers represent 75-95% quantiles, dots are data points outside of the 95% quantiles. Blue boxes = chorus present, gray boxes = chorus absent. Note that the chorus at GR01 is nearly only present with 0% lunar illumination, squeezing the box together at the bottom of the graph.

### Influence of environmental factors on chorus peak frequency

The choruses displayed clear peak frequencies on most nights, although some nights also contained one or more lower level choruses that deviated in peak frequency from the main chorus (Fig. S5). Initial exploration of the data indicated that seasonality and temperature were collinear. Peak frequency of the chorus varied with temperature, while chorus level varied with seasonality (Fig. 5). Therefore, we included temperature in our models for chorus peak frequency and month for our models on chorus level. The peak frequency of both Atlantic and plainfin midshipman choruses increased with increasing temperature (3-6 Hz per 1 °C, Table S2) at all but one site, MB02 (Fig. 6). MB02 also had less variability in water temperature and maintained cooler waters.

**Figure 5:**
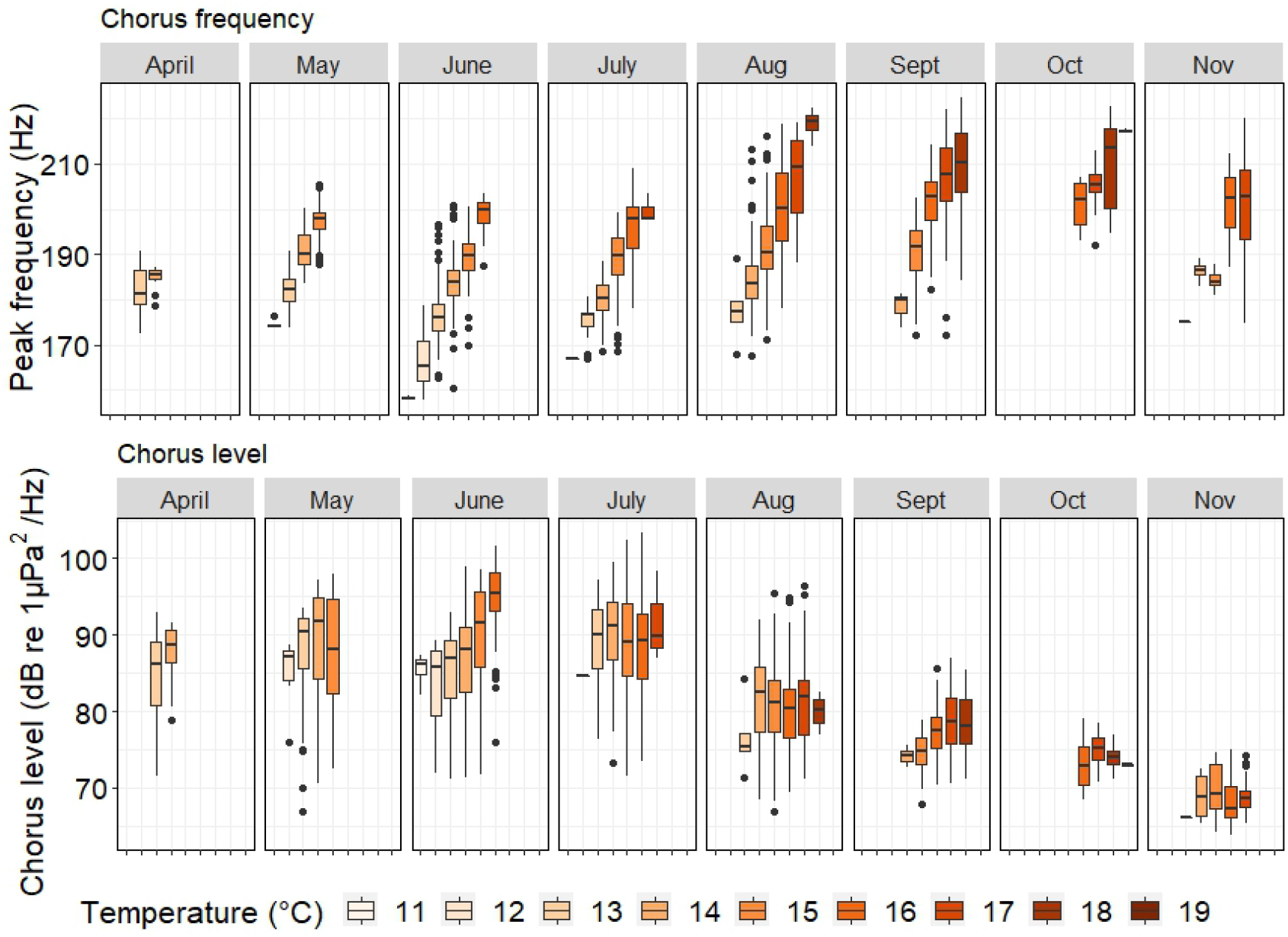
The peak frequency of plainfin midshipman chorusing increased with increasing temperature, but was not related to month of the year (top panel). In contrast, the chorus level changed over the year, peaking in July and then dropping off, but was not related to temperature (bottom panel). Cool to warm colors indicate increasing temperature, boxplots show 25 to 75 percentiles (box) with median (black bar) and 5 to 95 percentiles (whiskers). Dots represent data points that fell outside of the 95% percentile range. Results are split up per month and documented for site CI01.

**Figure 6:**
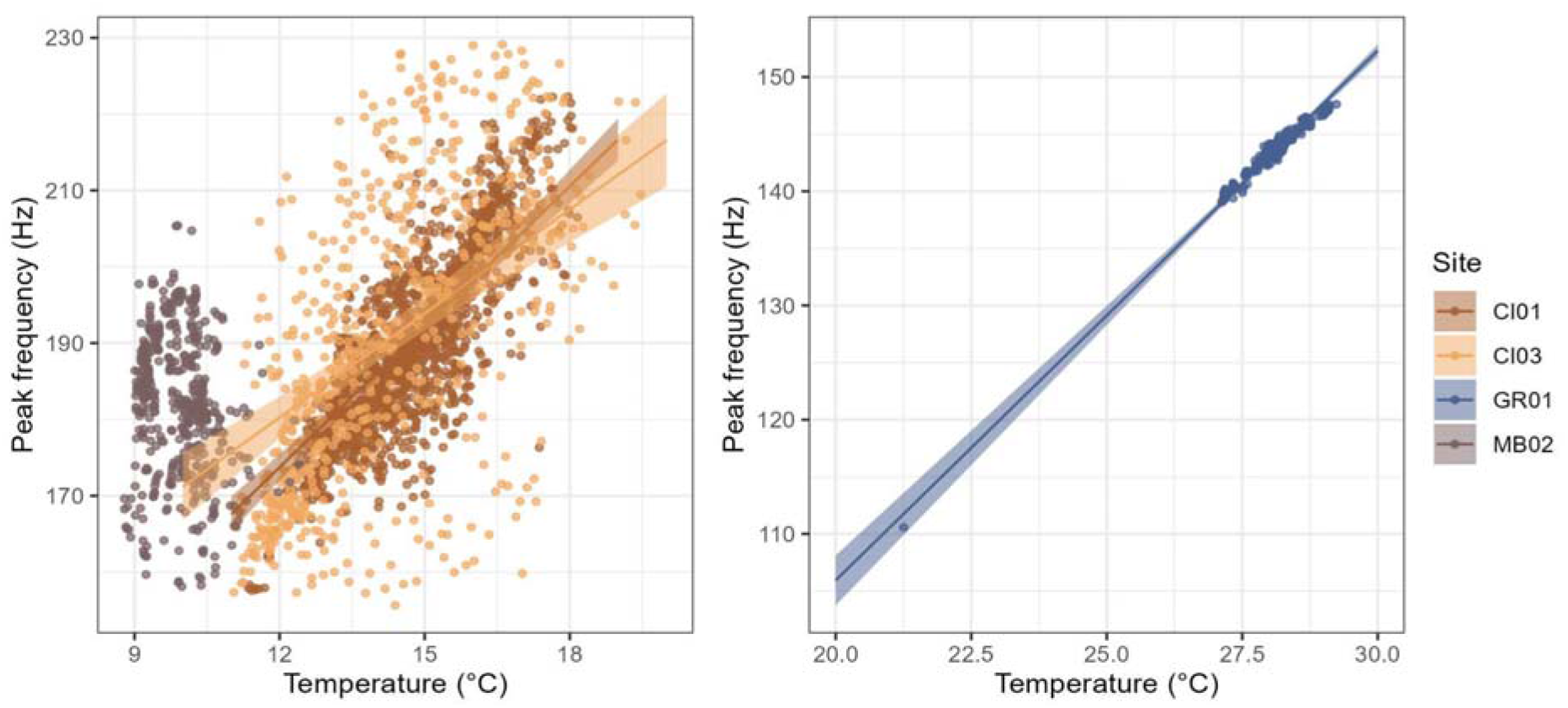
Peak frequency of midshipman chorus showed a positive relationship with water temperature. Correlations could be found at all sites except for the generally cooler sites in Monterey Bay (MB). Note that MB01, FK01, FK02 and FK03 were excluded from this analysis, due to the low presence of high SNR chorusing at those sites.

At CI01, CI03 and GR01, increasing lunar illumination was correlated to a slightly decreasing chorus peak frequency of <1 Hz per percent illumination (CI01: est = -0.017, p<0.0005; CI03: est = -0.25, p<0.05; GR01: est = -0.18, p<0.01, Table S2). Both at CI03 and GR01, this decrease was mostly visible at lower temperatures and could be related to overrepresentation of some temperatures in the data (CI03: est = 0.016, p<0.05, GR01: est = 0.0064, p<0.01). Noise levels did not correlate with chorus peak frequency at most sites. However, at CI03, chorus peak frequency slightly decreased with increasing noise levels (<1 Hz per 1 dB; est = -3.24, p<0.0005), but this was only the case at low temperatures (Fig. S4). An examination of the data distribution showed that high noise levels correlated with high wind speeds (Pearson correlation coefficient = 0.36, p<0.0001) and low temperatures in July.

### Influence of environmental factors on chorus SPL

Chorus SPL increased as the season progressed, peaking in the middle of the season in summer and then dropping off towards fall (Fig. 7). At three of the four sites that were included in this analysis (CI01, CI03, and MB02), higher lunar illumination was related to slightly lower chorus levels for some months (<1 dB per 1% illumination; Table S3), with the strongest effect at CI03. Ambient noise levels were correlated with increased chorus levels for some months at CI01 and CI03 (CI01: noise:month spline 4: estimate = 1.30, p<0.01; CI03: noise:month spline 2: estimate = 1.68, p<0.05), increasing average chorus levels with <2 dB (Table S3). At CI01, this correlation was visible at the end of the season, while at CI03, chorus levels only increased with noise in June and July. At MB02, there was a significant influence of noise during all months except June, but the data simulation suggested that results could only be trusted in June and July for this site (Fig. S3, second column). At GR01, there was a significant influence of noise in August (noise:month spline 2: estimate = -7.92, p<0.0001, Table S3).

**Figure 7:**
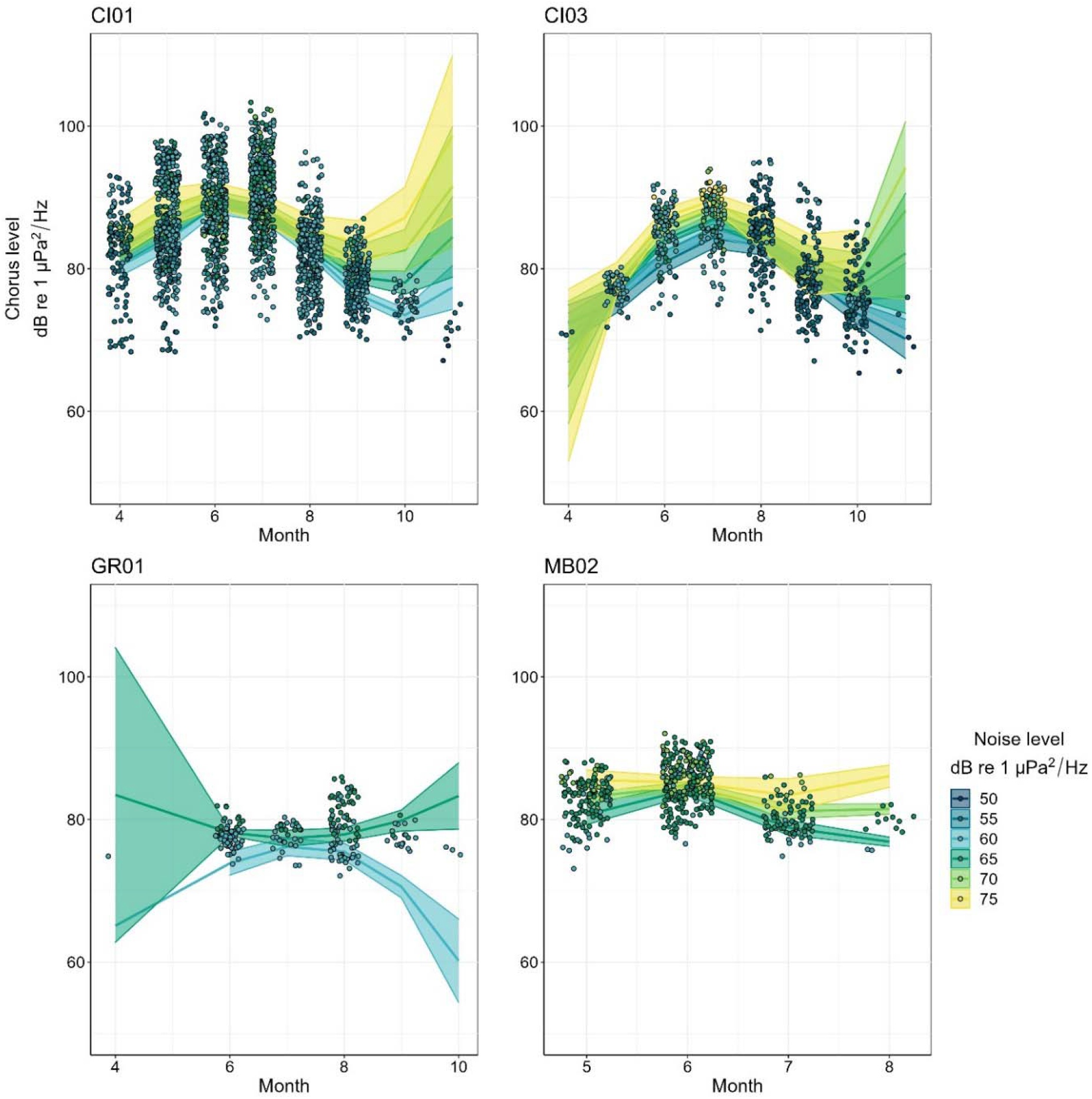
Chorus levels changed over the season, peaking around June-August, and showed slight increases with increasing noise levels for some months (color gradient in 5 dB bins; dots = data, lines = mean predicted chorus levels for selected noise levels representative of data, shaded areas = 95% CI). Chorus levels are shown per month, with some jitter in the data points to avoid overlap. Note that MB01, FK01 and FK02 were excluded from this analysis, due to the low presence of high SNR chorusing at those sites.

## DISCUSSION

In this study, we show that both plainfin midshipman and putative Atlantic midshipman changed their calling behavior in relation to environmental conditions. They predominantly called at night, at the shallow sites preferably when the moon was absent and showed peak calling in spring and early summer. Their chorus peak frequency increased as the water temperature increased. For both presumed Atlantic and plainfin midshipman chorus levels were only marginally affected by lunar illumination and were in some months influenced by ambient noise levels.

### Atlantic midshipman

The chorus of Atlantic midshipman has never been unambiguously described (Wall et al. 2013; Wall et al. 2014). Although we cannot be certain that the chorus we describe here is that of Atlantic midshipman, several facts point to that conclusion. First, based on the similarities of the chorus to that of plainfin midshipman, it is likely that the chorus is produced by a species of the genus *Porichthys*. Second, the only species of that genus native to the study region of the western North Atlantic is *Porichthys plectrodon*, or Atlantic midshipman (Collette 2002). Other species in that genus are found further south, from Honduras down to South America, making it unlikely that they would be encountered as far north as Gray’s Reef NMS. Thus, the chorus detected is likely produced by Atlantic midshipman, but future efforts are needed to confirm this presumption.

### Temporal patterns

Chorus presence was less likely during nights with high lunar illumination, for all but the two sites at Monterey Bay NMS. This difference in result between sites is likely related to decreased light penetration with water depth, as both Monterey Bay NMS sites were significantly deeper than the others (>80 m vs <30 m). Moon light is known to affect acoustic activity of many species, both aquatic and terrestrial (Dickerson et al. 2023). Increased light conditions from the moon facilitate multimodal communication, but also increase susceptibility to predation (Dickerson et al. 2023). This trade-off leads some fish species to call more close to full moon (Mccauley 2012; Borie-Mojica et al. 2022), while others restrict their calling to periods with little lunar illumination (Mooney et al. 2016; Parsons et al. 2016; Koenig et al. 2017; Kaplan et al. 2018). This can even vary within species. For instance, cusk eel calling was highest on new moon nights for animals living on sandy substrate (Mooney et al. 2016), while cusk eels living in eelgrass – with lower risk of visual predation – called most on full moon nights (Wirth and Warren 2020). Another driving factor could be the predation pressure on offspring. A reef fish species, the Atlantic goliath grouper (*Epinephelus itajara*), increased its spawning effort on new moon nights, when visually hunting egg predators were less abundant (Koenig et al. 2017).

### Influence of temperature

Positive relationships between water temperature and call peak frequency of fish have been found for multiple fish species, including plainfin midshipman (McIver et al. 2014; Ladich 2018). For midshipman fish, the contraction rate of the sonic muscle increases with increasing temperature ((Bass and Baker 1991; Feher et al. 1998), but review (Ladich 2018)), likely related to fish being predominantly cold blooded. When the surrounding water temperature increases, the muscle becomes warmer and is able to contract faster. However, previous studies were conducted on short time scales and therefore have not been able to disentangle the influence of seasonality and temperature on call peak frequency (Ladich 2018). The temperature variability across the midshipman chorusing season enabled us to show that midshipman chorus peak frequency is indeed mediated by temperature rather than seasonality and is conserved between species.

### Influence of noise

Midshipman fish slightly increased hourly chorus levels with higher noise levels. Measurement artifacts are a known risk of bioacoustics studies (Brumm et al. 2017) and need to be thoroughly considered before drawing conclusions about impact of noise on animal calling behavior. Since the noise levels included in the data were relatively low amplitude (50-80 dB re 1uPa/Hz) and measured chorus levels in the cleaned dataset were typically 80-90 dB re 1uPa/Hz, it could be that at those noise levels, there was no need for midshipman fish to significantly increase their chorus level to maintain an effective communication range. Indeed, chorus levels increased significantly only in months with relatively high noise levels, such as July at CI03. Perhaps the pressure to overcome anthropogenic noise was weak in the measured contexts because the chorusing peaks were at night, offset from peak vessel-associated human activity during the day.

Alternatively, it could be that the Lombard effect in midshipman fish is small, or that fish were not able to maintain increased chorus levels for an hour. A previous, experimental study on the Lombard effect in plainfin midshipman found an increase in midshipman call levels during noise of ∼90 dB, but the increase in call levels was small and mostly driven by an increased variation in call levels (Brown et al. 2021). This could indicate that not all individuals were able to increase their call levels. If midshipman fish truly have a limited capacity to increase their call levels with increasing noise, they are at risk of masking if noise levels increase more, leading to reductions in communication range with conspecifics (Erbe et al. 2016).

## Conclusion

Environmental factors influenced the calling behavior of both plainfin midshipman and Atlantic midshipman. In contrast to experimental settings, observational studies in the field often have to deal with various drivers that simultaneously influence the behavior of interest. Only temporally and spatially replicated studies can elucidate these patterns and truly come to an understanding of the impact of the environment on fish calling behavior. The current study shows the amount of variability that exists between locations, which can be related to variability in environmental conditions, differences in proximity to the recorded fish, or variability between populations. Even so, the strongest environmental drivers showed consistent patterns between recording sites and even between species.

## ACKNOWLEDGMENTS

The authors would like to acknowledge the vessel crews and coordinators of *R/V Fulmar*, R4107, *R/V Shearwater, R/V SharkCat, R/V Sam Gray, R/V Manta*, and FL FWC research fleet for their assistance with data collection, particularly Jackie Buhl, Zac Montgomery, Marshall Stein, Ryan Freedman, Rayon Carruthers, Clyde Terrell, Brian Yannutz, Jean de Marignac, Dave Lott, Kristin Raja, Nick DeProspero, Kimberly Roberson, Alison Soss, Lonny Anderson, Danielle Morley, and Jessica Keller. We thank Eduardo Ruiz-Morales for helping with the data analysis. This work was completed as part of the SanctSound project, which is a collaboration between NOAA and the U.S. Navy to better understand underwater sound within the National Marine Sanctuary System.

