## Supplementary material for "Impact of environmental factors on calling behavior in midshipman fish across ocean basins"

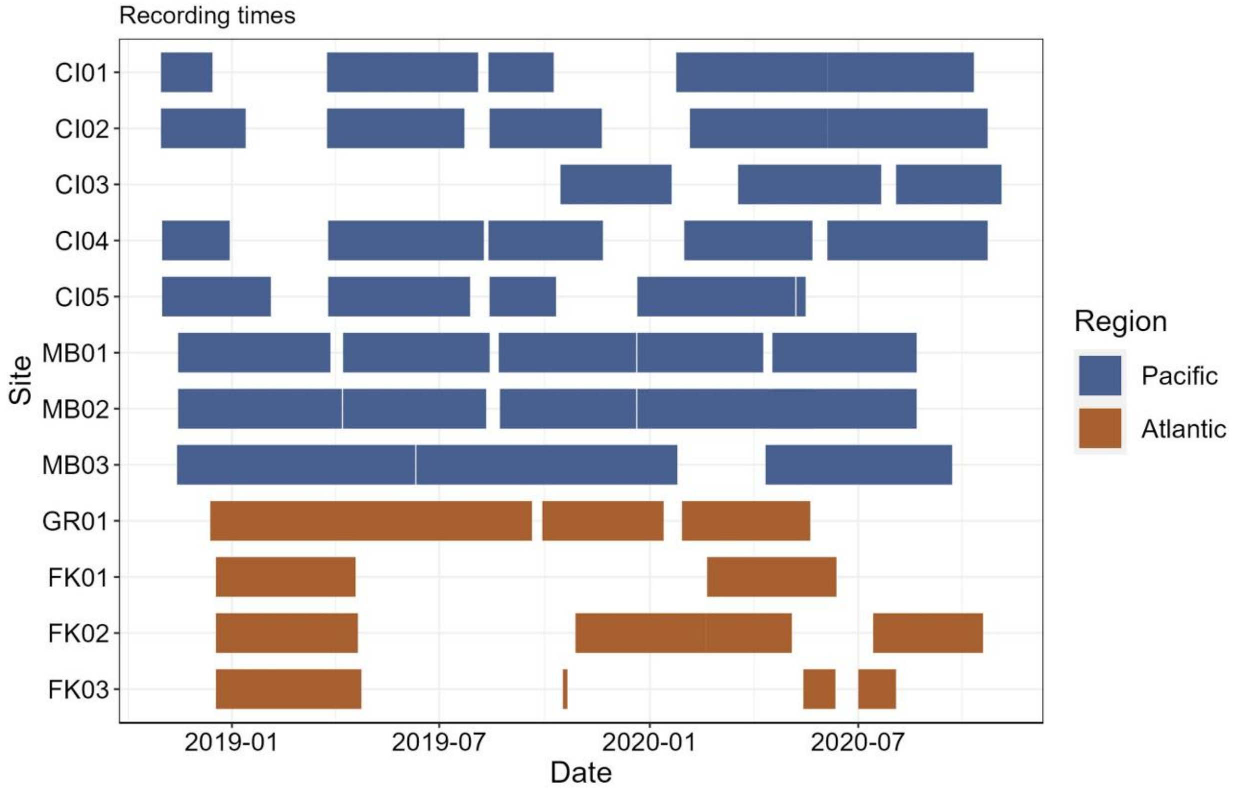


*Figure S1: Acoustic recording period ranged from November 2018 to August 2020, with gaps when the recorder had to be switched out or equipment malfunctioned. Top eight rows represent Pacific sites and bottom four rows represent Atlantic sites.*

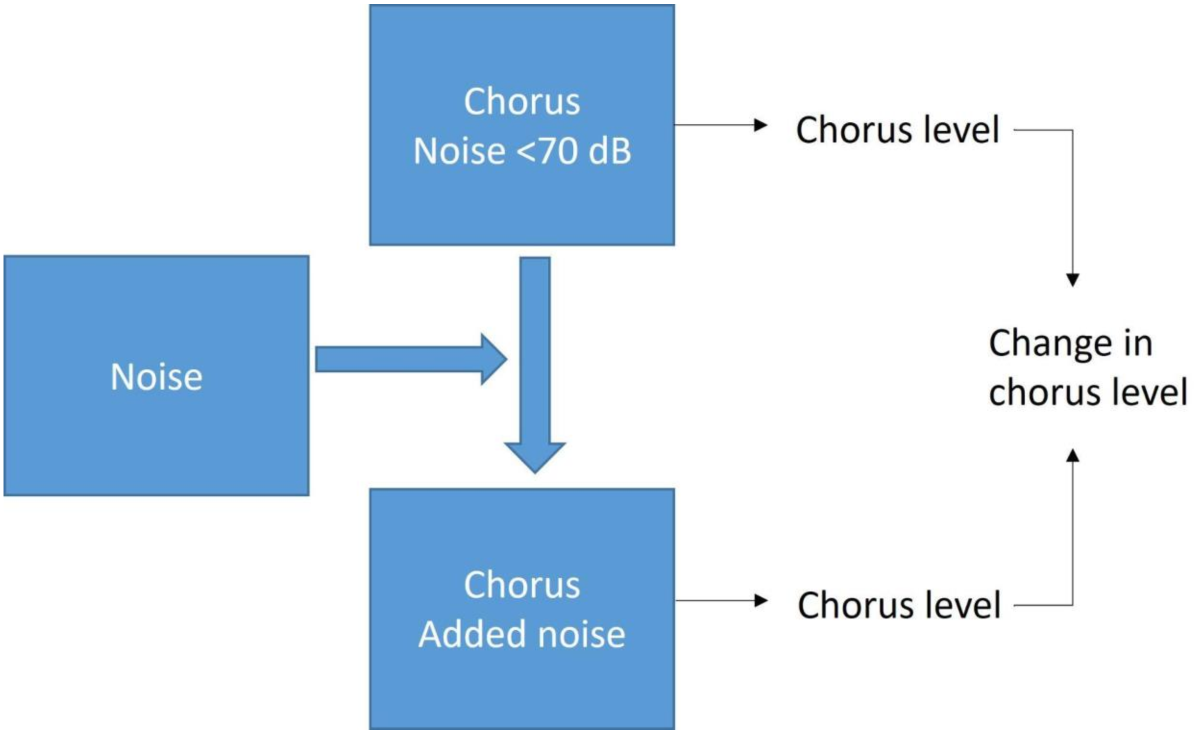


*Figure S2: Schematic representation of creating and analyzing the simulated dataset. Hours of recordings with chorus presence and noise levels below 70 dB were selected, to which noise from a different section of the recording, which did not include chorusing, was added on a minute-by-minute basis. Chorus levels were measured for every minute before and after adding noise. These measurements were then averaged to hourly medians, and subtracted from each other, to come to the change in chorus level because of added noise. A value of 2 dB or higher in the changed chorus level was considered to represent a measurement that was affected by the added noise.*

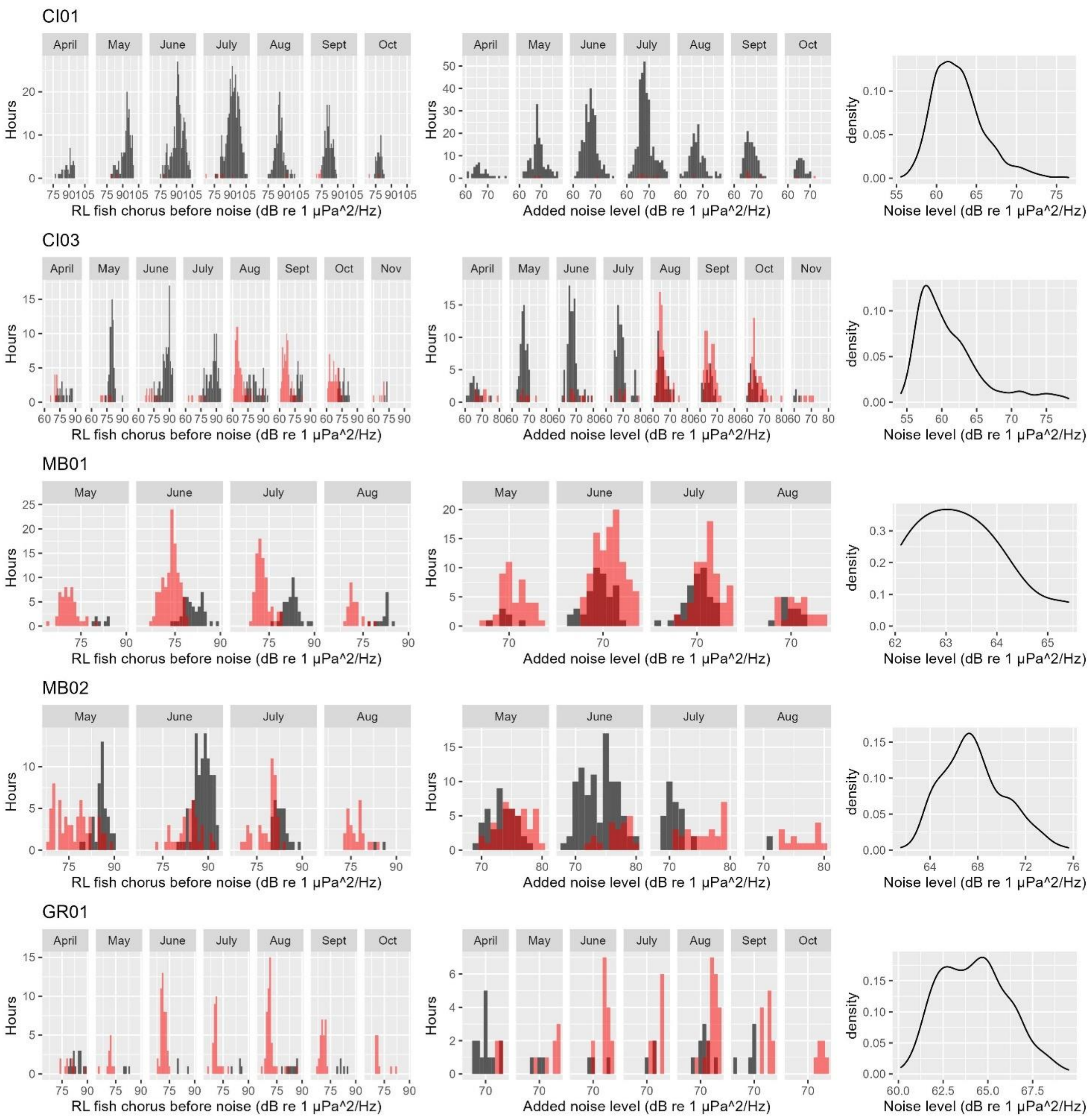


*Figure S3: Simulation for the effect of increasing noise levels on hourly median chorus level measurements. Rows show different sites. The first two columns show the distribution of the simulated data (only data of SNR>10 on minute level before taking hourly median): left: received level of the fish chorus before adding noise; middle: the level of the added noise. As a comparison, the right column shows the noise level distribution in the cleaned data set. Histograms show count of hourly chorus level measurements in 1 dB bins per month. Black bars = measurements that changed <2 dB after adding noise to the recording, red bars = measurements that changed >2 dB after adding noise to the recording. Density plots show the relative contribution of each hourly noise level to the data set of one site.*

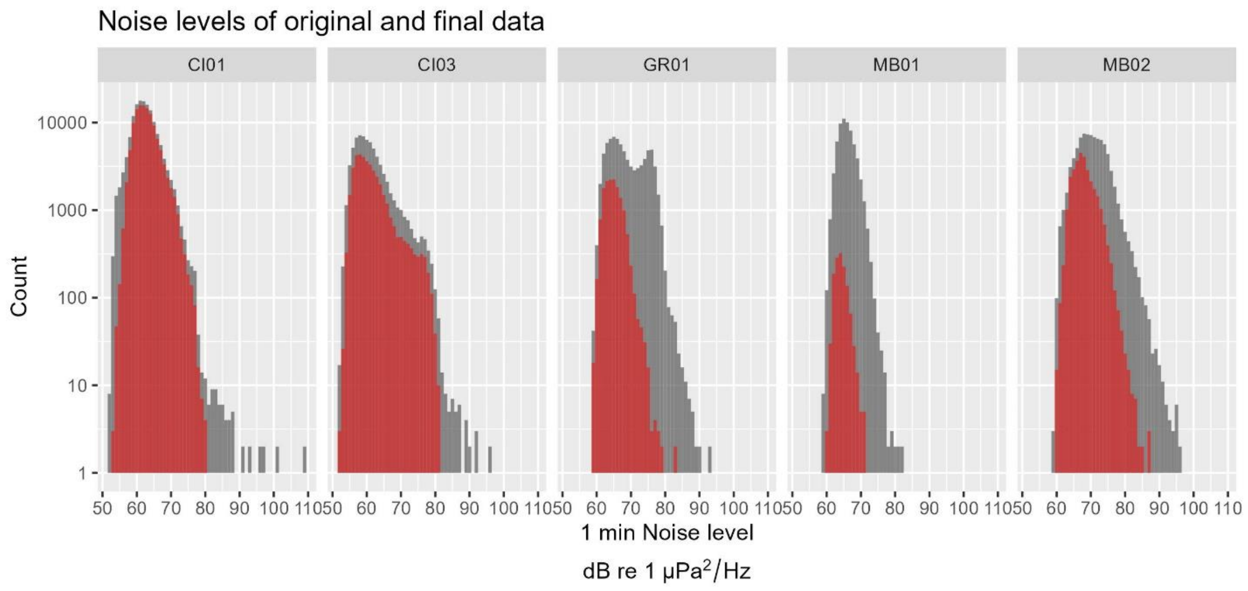


*Figure S4: Distribution of measured noise levels in 1 min bins of the original (gray bars) and final data set (red bars). Note that the y-axis is on a 10-log scale, to ensure visibility of all bars.*

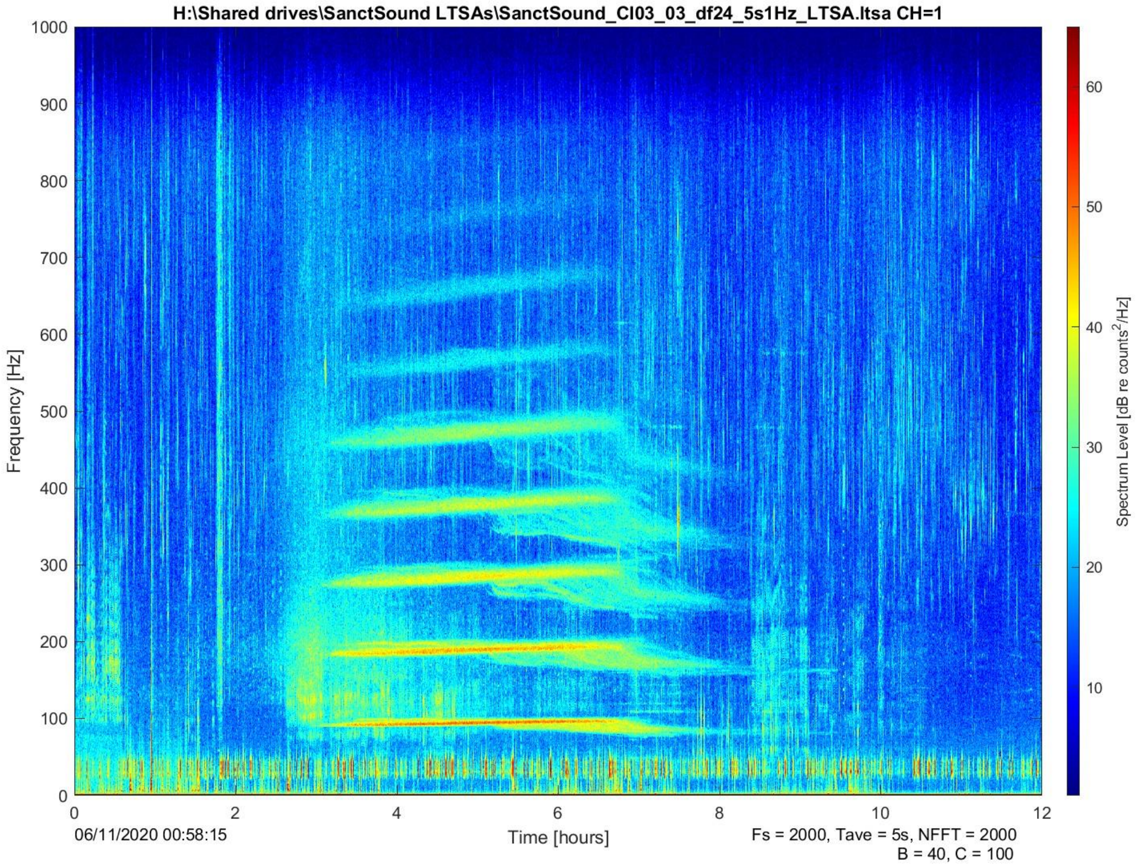


*Figure S5: Pacific midshipman chorus from CI03 on June 11, 2020. Midshipman choruses typically consisted of a strong peak frequency, which was likely made up of the calls of several individuals calling at the same frequency. However, calls of animals deviating in frequency could be visible as well, as shown here in hour 6-8.*

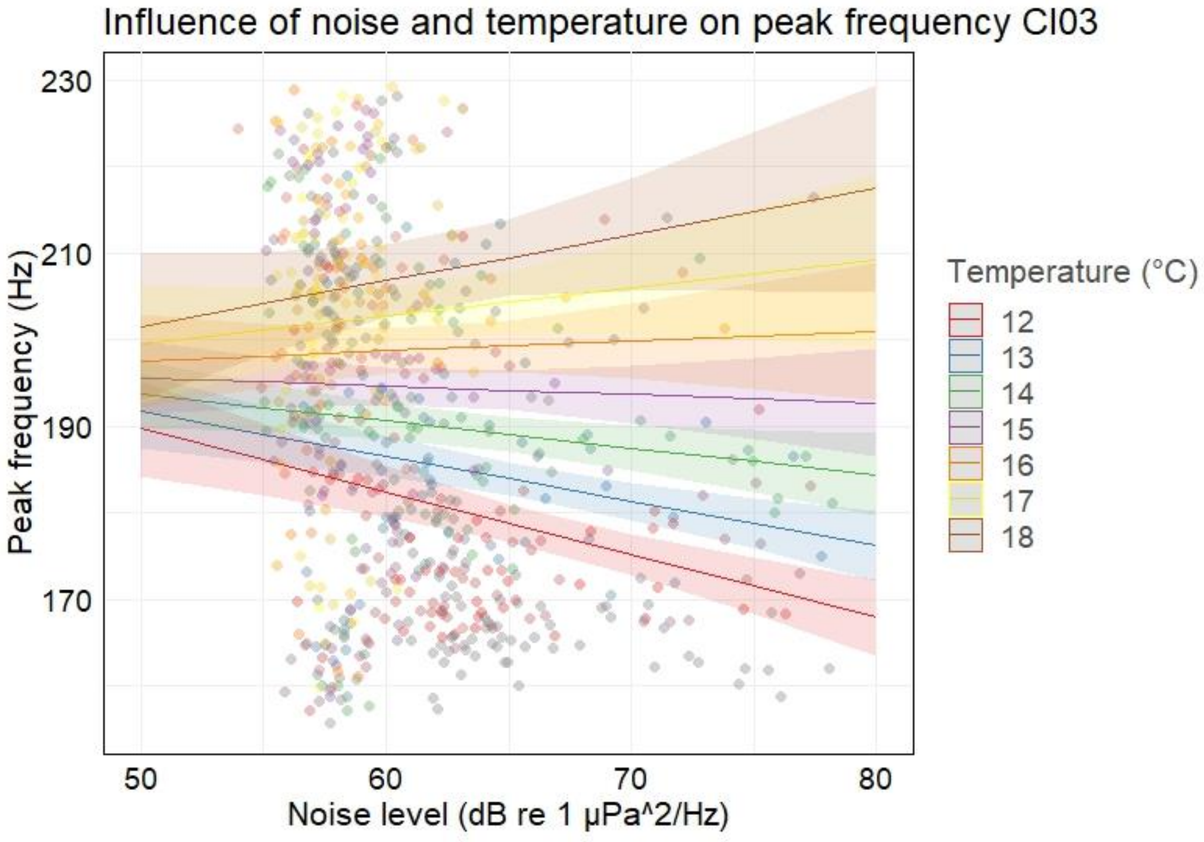


*Figure S6: Higher noise levels were correlated with lower peak frequencies at CI03, but this was mostly the case for low temperatures. Dots represent measured values, lines and shaded areas represent modeled estimates and confidence intervals, respectively. Similar colors show measurements at the same temperature.*

*Table S1: Modeling results of the effect of lunar illumination per site on hourly chorus presence. Values were calculated using a generalized linear model with binomial distribution and site MB02 in the intercept. Explanatory variables were lunar illumination, site, and the interaction between lunar illumination and site.*

| Coefficients | Estimate | SE | Z | p |
|---|---|---|---|---|
| *Intercept* | -1.71 | | | <0.0001 |
| Lunar illumination | -0.0005 | 0.0014 | 0.36 | 0.72 |
| *Site* | | | | |
| CI01 | 1.08 | 0.080 | 13.58 | <0.0001 |
| CI03 | 0.87 | 0.091 | 9.58 | <0.0001 |
| MB01 | -3.28 | 0.44 | -7.39 | <0.0001 |
| GR01 | -0.62 | 0.11 | -5.53 | <0.0001 |
| FK01 | -0.77 | 0.15 | -5.13 | <0.0001 |
| FK02 | -0.27 | 0.10 | -2.70 | <0.01 |
| FK03 | -0.013 | 0.17 | -0.073 | 0.94 |
| *Site:lunar* | | | | |
| CI01:lunar | -0.0051 | 0.0016 | -3.23 | <0.005 |
| CI03:lunar | -0.019 | 0.0021 | -8.89 | <0.0001 |
| MB01:lunar | 0.0070 | 0.0074 | 0.95 | 0.34 |
| GR01:lunar | -0.018 | 0.0030 | -6.02 | <0.0001 |
| FK01:lunar | -0.020 | 0.0049 | -4.06 | <0.0001 |
| FK02:lunar | -0.089 | 0.0022 | -3.99 | <0.0001 |
| FK03:lunar | -0.015 | 0.0055 | -2.71 | <0.01 |

*Table S2: Modeling results of the effect of temperature, lunar illumination and noise level on the peak frequency of midshipman chorus. Each site was modeled separately using Generalized Estimating Equations with a Gaussian distribution, a first-order autocorrelation with day as a blocking unit and robust standard error. Only the explanatory variables that were retained in the final models are shown here.*

| Site | Coefficients | Estimate | SE | Wald | P |
|---|---|---|---|---|---|
| CI01 | *Intercept* | 100.26 | | | <0.0001 |
| | Temperature | 6.13 | 0.31 | 390.1 | <0.0001 |
| | Lunar illumination | -0.02 | 0.0048 | 12.4 | <0.0005 |
| CI03 | *Intercept* | 332.10 | | | <0.0001 |
| | Temperature | -8.76 | 4.11 | 4.55 | <0.05 |
| | Lunar illumination | -0.25 | 0.10 | 6.18 | <0.05 |
| | Noise | -3.24 | 0.86 | 14.22 | <0.0005 |
| | Temp:lunar | 0.016 | 0.0073 | 4.96 | <0.05 |
| | Temp:noise | 0.21 | 0.064 | 10.84 | <0.001 |
| MB01 | *Intercept* | 168.88 | | | |
| | Temperature | 1.17 | 0.94 | 1.55 | 0.21 |
| MB02 | *Intercept* | 192.07 | | | |
| | Temperature | -1.09 | 1.31 | 0.69 | 0.4 |
| GR01 | *Intercept* | 17.59 | | | <0.0001 |
| | Temperature | 4.48 | 0.13 | 1260.47 | <0.0001 |
| | Lunar illumination | -0.18 | 0.066 | 7.59 | <0.01 |
| | Temp:lunar | 0.0064 | 0.0023 | 7.47 | <0.01 |

*Table S3: Modeling results of the effect of month, lunar illumination and noise level on midshipman chorus levels. Each site was modeled separately using Generalized Estimating Equations with a Gaussian distribution, a first-order autocorrelation with day as a blocking unit and robust standard error. Month was included as an m-spline with four knots. Only the explanatory variables that were retained in the final models are shown here.*

| Site | Coefficient | Estimate | SE | Wald | P |
|---|---|---|---|---|---|
| CI01 | *Intercept* | 76.23 | | | <0.0001 |
| | Noise | 0.12 | 0.086 | 1.84 | 0.18 |
| | Month (spline1) | -35.07 | 26.63 | 1.67 | 0.19 |
| | Month (spline2) | 94.74 | 44.13 | 4.61 | <0.05 |
| | Month (spline3) | -79.54 | 46.96 | 2.87 | 0.09 |
| | Month (spline4) | -83.80 | 29.13 | 8.28 | <0.005 |
| | Moon | -0.044 | 0.011 | 17.47 | <0.0001 |
| | Noise:monthS1 | 0.55 | 0.41 | 1.79 | 0.18 |
| | Noise:monthS2 | -0.97 | 0.69 | 2.01 | 0.16 |
| | Noise:monthS3 | 0.75 | 0.76 | 0.98 | 0.32 |
| | Noise:monthS4 | 1.30 | 0.51 | 6.73 | <0.01 |
| | MonthS1:moon | -0.015 | 0.035 | 0.18 | 0.67 |
| | MonthS2:moon | 0.13 | 0.054 | 6.01 | <0.05 |
| | MonthS3:moon | 0.033 | 0.061 | 0.29 | 0.59 |
| | MonthS4:moon | 0.069 | 0.040 | 3.05 | 0.08 |
| CI03 | *Intercept* | 96.85 | | | <0.0001 |
| | Noise | -0.44 | 0.39 | 1.32 | 0.25 |
| | Month (spline1) | -69.83 | 54.71 | 1.63 | 0.21 |
| | Month (spline2) | -50.57 | 42.67 | 1.40 | 0.24 |
| | Month (spline3) | 13.08 | 71.02 | 0.03 | 0.85 |
| | Month (spline4) | -30.47 | 36.75 | 0.69 | 0.41 |
| | Moon | 0.013 | 0.044 | 0.09 | 0.76 |

| | | | | | |
|---|---|---|---|---|---|
| | Noise:monthS1 | 1.32 | 0.94 | 1.95 | 0.16 |
| | Noise:monthS2 | 1.68 | 0.68 | 6.13 | <0.05 |
| | Noise:monthS3 | -0.17 | 1.23 | 0.02 | 0.89 |
| | Noise:monthS4 | 0.64 | 0.63 | 1.05 | 0.30 |
| | MonthS1:moon | 0.0084 | 0.15 | 0.00 | 0.96 |
| | MonthS2:moon | -0.20 | 0.065 | 9.31 | <0.005 |
| | MonthS3:moon | 0.0040 | 0.13 | 0.00 | 0.98 |
| | MonthS4:moon | -0.15 | 0.053 | 8.05 | <0.005 |
| MB02 | *Intercept* | 50.06 | | | <0.0001 |
| | Noise | 0.48 | 0.13 | 14.21 | <0.0005 |
| | Month (spline1) | 55.33 | 18.96 | 8.51 | <0.005 |
| | Month (spline2) | -13.65 | 23.87 | 0.33 | 0.57 |
| | Month (spline3) | -25.00 | 8.48 | 8.70 | <0.005 |
| | Moon | -0.015 | 0.0063 | 5.55 | <0.05 |
| | Noise:monthS1 | -0.71 | 0.27 | 6.90 | <0.01 |
| | Noise:monthS2 | 0.11 | 0.35 | 0.10 | 0.75 |
| | Noise:monthS3 | 0.34 | 0.12 | 7.62 | <0.01 |
| | monthS1:moon | 0.0011 | 0.013 | 0.01 | 0.93 |
| | monthS2:moon | 0.029 | 0.011 | 6.68 | <0.01 |
| | monthS3:moon | 0.0083 | 0.0051 | 2.67 | 0.10 |
| GR01 | *Intercept* | -176.41 | | | 0.29 |
| | Noise | 4.06 | 2.71 | 2.24 | 0.13 |
| | Month (spline1) | 421.59 | 479.82 | 0.77 | 0.38 |
| | Month (spline2) | 500.82 | 124.74 | 16.12 | <0.0001 |
| | Month (spline3) | 45.59 | 337.85 | 0.02 | 0.89 |
| | Noise:monthS1 | -6.85 | 7.76 | 0.78 | 0.38 |

| | | | | |
|---|---|---|---|---|
| Noise:monthS2 | -7.92 | 2.01 | 15.52 | <0.0001 |
| Noise:monthS3 | -0.85 | 5.43 | 0.02 | 0.88 |
